# Cadherin Preserves Cohesion Across Involuting Tissues During *C. elegans* Neurulation

**DOI:** 10.1101/2020.05.05.079301

**Authors:** Kristopher Barnes, Li Fan, Mark W. Moyle, Christopher Brittin, Yichi Xu, Daniel Colón-Ramos, Anthony Santella, Zhirong Bao

## Abstract

The internalization of the central nervous system, termed neurulation in vertebrates, is a critical step in embryogenesis. Open questions remain as to how force propels coordinated tissue movement during the process, and little is known as to how internalization happens in invertebrates. We show that in *C. elegans* morphogenesis, apical constriction in the retracting pharynx drives involution of the adjacent neuroectoderm. Localized HMR-1/Cadherin mediates the inter-tissue attachment, as well as within the neuroectoderm to maintain intratissue cohesion. Our results demonstrate that localized HMR-1 is capable of mediating embryo wide reorganization driven by a centrally located force generator, and indicate a non-canonical use of Cadherin on the basal side of an epithelium that may apply to vertebrate neurulation. Additionally, we highlight shared morphology and gene expression in tissues driving involution, which suggests that neuroectoderm involution in *C. elegans* is potentially homologous with vertebrate neurulation and thus may help elucidate the evolutionary origin of the brain.

## Introduction

### Nervous System Centralization

Neurulation is the process which establishes the centralized position of the developing nervous system in vertebrates via involution of the neuroectoderm, with failure resulting in the Neural Tube Defect category of diseases (Gilbert 2000). During involution, the neuroectoderm layer bends inwards at its center until it is fully internalized. This physical change is part of a coordinated tissue movement, which requires interaction between the neuroectoderm and adjacent tissues. The floor plate, which generates the pulling force and is required for involution, is derived from a separate tissue despite ending up in the same epithelial layer as the neuroectoderm (Smith 1997). The notochord, which is well known for its Sonic Hedgehog signals to pattern neuron subtypes in the neural tube, also cohesively attaches to the floor plate and is necessary to stabilize the involuting ectoderm during the process (Smith 1997). The attached epidermis is also in-part pulled by the neuroectoderm; failure of the epidermis to close during neurulation contribute to Neural Tube Defect diseases (Gordon 1985). Which factors regulate the interaction between adjacent tissues is a particularly understudied aspect of neurulation.

Apical constriction produces the force required to reshape the neuroectoderm (Sawyer et al 2010) and is the strongest in the floor plate. During apical constriction force is generated via contraction of an apical circumferential actomyosin belt within cells, with tight junctions and adherens junctions anchoring the belt between cells and enabling the transmittance of force across the cell layer. Cadherin/catenin complexes at lateral adherens junctions are necessary to hold the epithelial layers together in addition to their role in enabling constriction (Ilina and Fiedl 2009). In addition, planer cell polarity (PCP) and convergent extension are important in elongation across the A-P axis (Williams et al 2014). Still, there remains much to be understood regarding the complex way that force production and tissue cohesion interact to enable neuroectoderm involution during neurulation.

### *C. elegans* Nervous System Formation

The *C. elegans* nervous system was first mapped in its entirety by John White (White et al 1986), and the full and invariant cell lineage was determined by John Sulston (Sulston et al 1983). The central nervous system is composed of the nerve ring, the main neuropil composed of 181 axons, as well as the ventral nerve cord (VNC) (White et al 1986). Most neurons in the embryo are born between 300-320 minutes post-fertilization (mpf) and proceed to internalize and move nearer to the midline of the embryo (Harrel and Goldstein 2011), a process which has not been characterized in depth. Neurons subsequently begin projecting axons around early comma stage (∼360 mpf) and proceed to form a visible ring within an hour (Santella et al 2015, Rapti et al 2017, Moyle et al 2020). During ventral cleft closure (Chisholm and Hardin 2005), the initial movement of the VNC neurons to the midline and subsequent reorganization via PCP mediated cell intercalation and convergent extension have been characterized (George et al 1998, Shah et al 2017). However, the process behind internalization of the neurons in the head, which will go on to form the nerve ring, has not been characterized.

These developmental events in the nervous system occur simultaneously with major morphogenesis events in non-neuronal tissues including the pharynx and skin. The head nervous system develops alongside the pharynx (an organ unrelated to the vertebrate pharynx), and the nerve ring ultimately encircles it. Signaling from the pharynx during morphogenesis regulates the anterior-posterior placement of the nerve ring (Kennerdell et al 2009). The bilayer pharyngeal primordium, located at the center of the nascent head through gastrulation (Harrel and Goldstein 2011, Pohl et al 2012), retracts into a bulb due to apical constriction during this time period (Santella et al 2010, Rasmussen et al 2012). This retraction is a major event in the course of head morphogenesis, but its link to the formation of the rest of the tissues in the head including that of the nervous system has not been studied.

The skin forms on the dorsal exterior of the embryo during bean stage and subsequently extends to close and seal at the anterior and ventral midline. The ventral neuroblasts have been shown to be required for proper ventral closure (George et al 1998), with skin crawling over neuron substrates (Wernike et al 2016). In the head, skin closure requires the Cadherin orthologue *hmr-1* (Costa et al 1998), but the morphogenesis process is less characterized.

In this paper, we connect the morphogenesis of the pharynx and skin to the internalizing movement of the neurons. We show that this movement involves the involution of a cohesive neuroectoderm layer driven by attachment to the retracting pharynx in a pattern with striking similarity to vertebrate neurulation, and we characterize the role of HMR-1 in establishing inter-tissue attachment and maintaining intra-tissue cohesion over the course of head formation.

## Results

### Nervous System Involution is a Coordinated Process Between the Pharynx and the Neuroectoderm

In order to characterize the process by which the nervous system internalizes in *C. elegans*, we examine head morphogenesis during the 60-minute time window between the terminal division of the neurons and initial axon outgrowth. We first examine cell movements through WormGUIDES, a 4D atlas of *C. elegans* embryogenesis that tracks the position and lineage identity of every nucleus at every minute from the 4-cell stage to the one-and-half fold stage, about 2 hours after terminal division of neurons (Santella et al 2015). The *C. elegans* neuroectoderm begins on the exterior of the embryo during early bean stage (∼300 mpf). The pharyngeal cells have formed a two-sheet structure in the center of the head after invaginating during gastrulation. Head neurons envelope the pharynx at this stage (∼300 mpf, Figure 1a first timepoint). One hour later the pharynx has contracted into a bulb, and the neuroectoderm has moved to the anterior and ventral midline (Figure 1a second timepoint, Figure 1 Movie 1, Figure 1 Movie 2), and effectively internalized.

**Figure 1:**
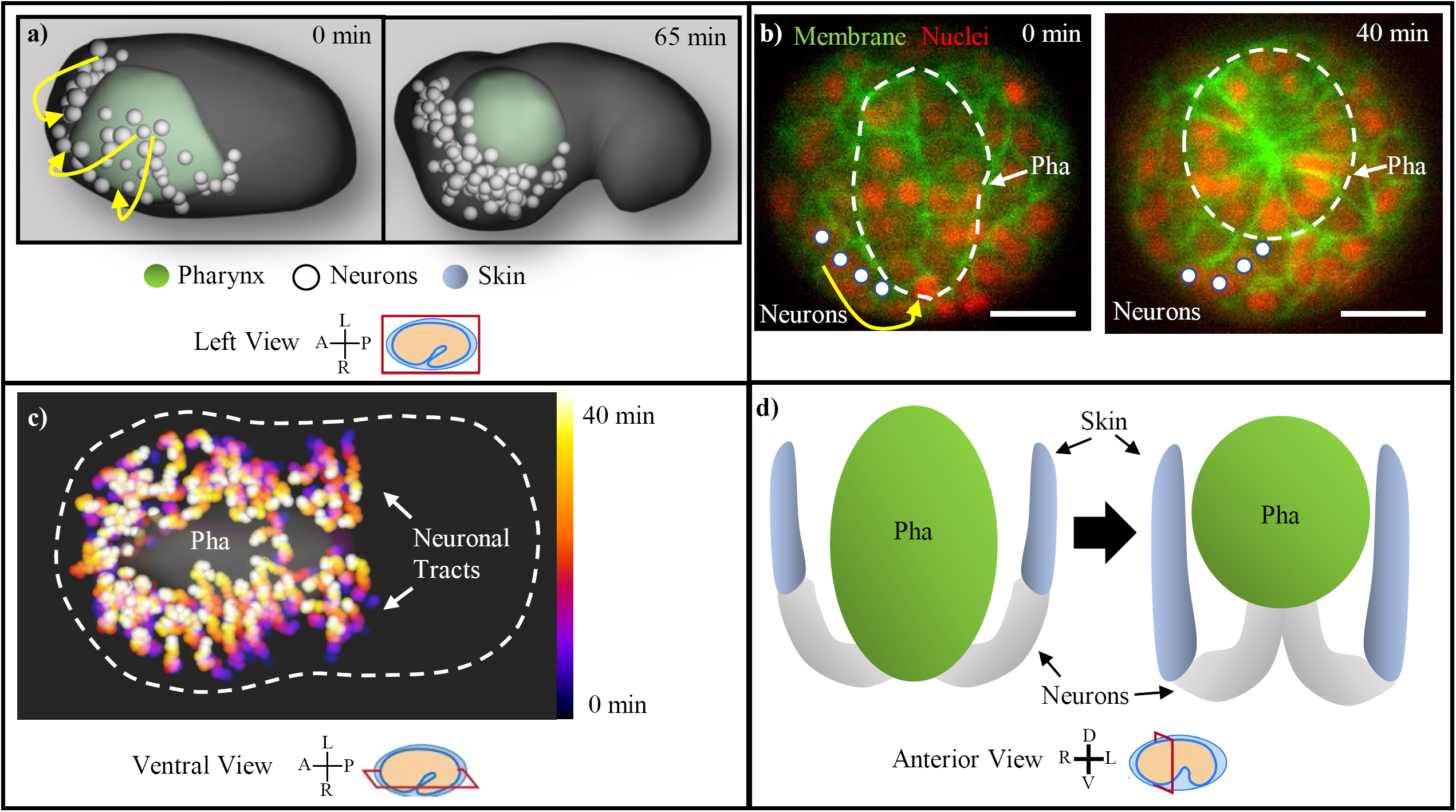
*C. elegans* Nervous System Centralization Consists of the Involution of the Neuroectoderm with the Retracting Pharynx. **(a)** Head neurons (white) are observed enveloping the pharynx (green) as visualized in the WormGUIDES app. Yellow arrows indicate the circumferential path of the involuting neurons. Axis compass and view plane are displayed in this format throughout all figure panels. **(b)** Neuron chain (marked by white circles) during involution, shown in *unc33p::PH::GFP* expressing embryos (transverse plane at mid-pharynx, marked by white dashed circle). Yellow arrow indicates path of involuting neurons. Red channel is cell nuclei. Scale bars are 10µm. **(c)** Temporal max projection of involuting neurons on WormGUIDES, with chains progressing over 40 minutes from black/purple to yellow/white. White dashed line is the embryo outline. **(d)** Model showing the pharynx (green), neuron chain (white), and skin (blue) before and after pharynx retraction and involution.

To better examine cell shapes and potential cohesive relationship among cells during this process, we conduct 3D, time-lapse imaging with a pan membrane marker (*unc-33p::PH::GFP*) and a ubiquitous nuclear marker (histone::mCherry) to track the cell lineage (Bao et al 2006, Santella et al 2014, Katzman et al 2018). In the transverse plane near the middle of the pharynx(Figure 1b), apical constriction of the two-sheet pharynx (Figure 1b, dashed circle), the inward/dorsal retraction of the pharynx, and the coordinated circumferential movement of neurons (Figure 1b, white dots and arrow) following the retracting pharynx are evident. Persistent cell contact is maintained between the neurons interfacing with the pharynx, as well as among the chain of involuting neurons. This coordinated movement occurs along the anterioposterior extent of the head. Visualized from the ventral side of the embryo (Figure 1c), movement trajectories of individual neurons (temporal max projections) show largely parallel tracts, which together with the persistent cell contacts observed above indicate tissue cohesion in the neuroectoderm and the pharynx during this movement (Figure 1c). Essentially all head neurons participate in this movement except for the amphid. During this process, the skin epithelium also moves anteriorly, with persistent cell contract between its leading edge and the trailing edge of the involuting neurons, before it eventually moves over the neurons to encase the head (Figure 1 supplement 1).

To explain the coordinated tissue movement and apparent tissue cohesion (Figure 1d), we propose a model in which retraction of the pharynx propels involution of the attached neuroectoderm. We examine this model below.

### Involution of the Head Neurons Requires *hmr-1*/Cadherin

Because *hmr-1/Cadherin* plays an important role in cell adhesion and is known to be widely expressed at this stage of embryogenesis (Achilleos et al 2010), we ask if we can perturb neuroectoderm involution by inducing *hmr-1* loss of function. To this end, we examine mutants homozygous for the *zu248* allele (Costa et al 1998) and use a *cnd1p::PH::RFP* marker to label a set of bilateral ventral neurons (Shah et al 2017) to measure their movements trajectories with live imaging. In the WT, these neurons migrate 25 microns over 60 min to meet at the midline (12/12). In *hmr-1* mutants, 75% of the embryos (9/12) fail to do so at the completion of pharynx retraction (Figure 2a), with an average of 28 microns separation between the bilateral neurons versus 0 microns for all WT (Figure 2b). 25% (3/12) of *hmr-1* embryos successfully involute but arrest soon after as neurons appear to detach at the midline. We conclude that *hmr-1* is required for successful involution of the head neuroectoderm.

**Figure 2:**
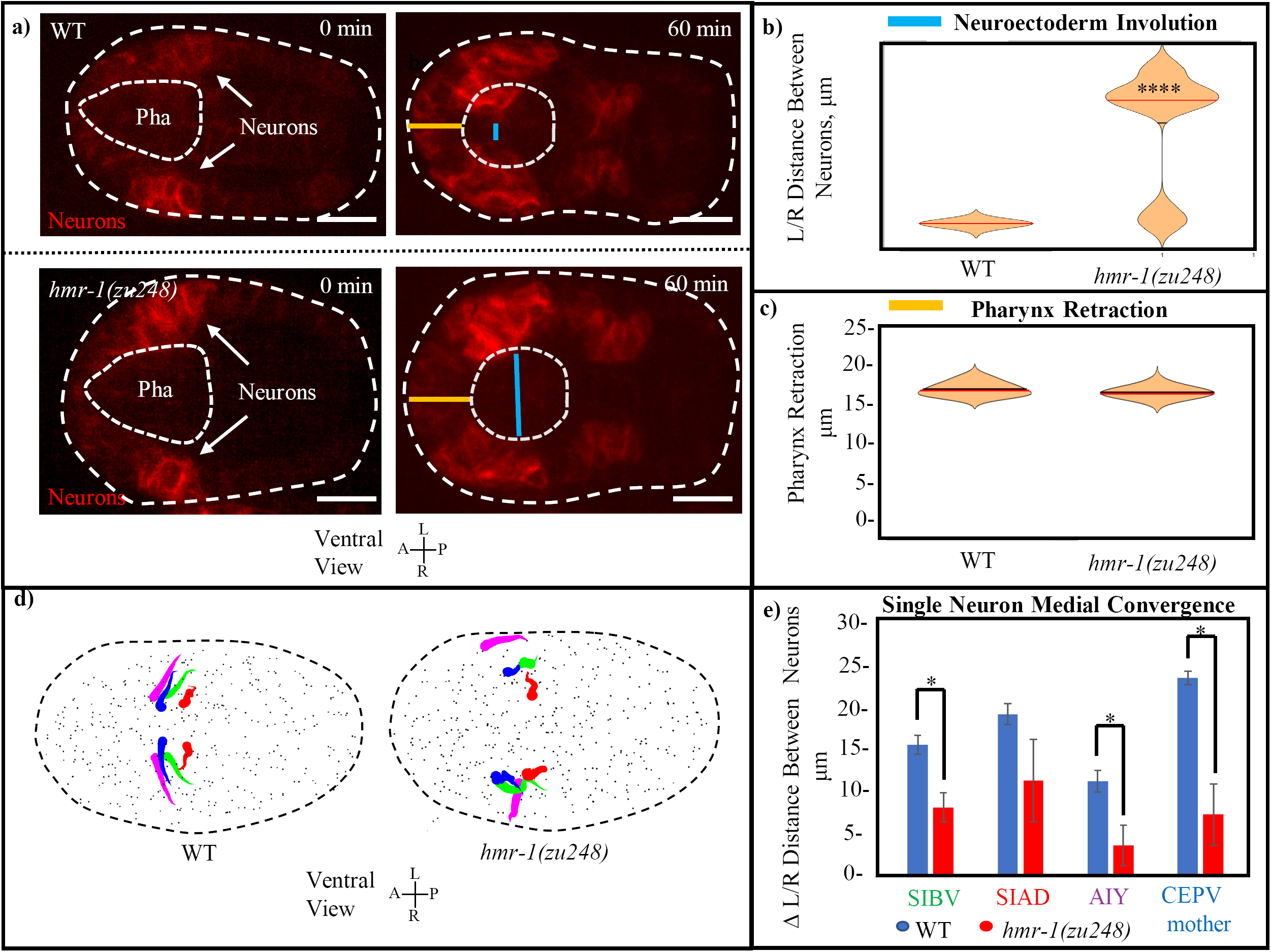
Involution of the *C. elegans* Head Neurons Requires HMR-1. **(a)** Ventral neurons in *cnd1p::PH::RFP* expressing embryos move towards the midline during involution. Distance between left/right leading edges (blue line) is visibly increased in *hmr-1(zu248)* mutants. Outer dashed line is embryo outline, inner dashed line is pharynx. Distance of pharynx retraction (orange line) is conserved. Scale bars are 10µm. **(b)** Involution of the neuroectoderm is measured as the distance between left and right side neurons in (n=10) *hmr-1(zu248)* embryos versus in (n=10) WT embryos (measurement of the blue line from panel a). Measurement is performed at the timepoint where the line is the shortest. Median is marked by red, significance was calculated with a 1 tailed student’s T-test. For all figures, *=p<0.05, and ****=p<0.001. **(c)** Pharynx retraction in WT vs *hmr-1(zu248)* embryos. Measurement is of length from anterior pole of embryo to anterior pole of pharynx (orange line in a). n=6 embryos (WT) and n=8 embryos *(hmr-1* mutant). Red line in plot marks median. **(d)** Motion paths of select neurons near the leading edge of involuting tissue in WT and *hmr-1(zu248)* embryos. Tadpole shape represents progression from early timepoints (tail) to late timepoints (head). Colors of neuron names in d) are the same as the color of the equivalent motion paths (left and right). Dashed line is embryo outline, grey dots are other cells. **(e)** Quantification of movement trajectories from c), measuring distance traveled by L/R partners towards each other. Significance across n=3 embryos was calculated with a 1 tailed student’s T-test.

To rule out the possibility that the lack of neuroectoderm involution is due to a lack of pharynx retraction, where HMR-1 is known to localize to its apical side (Sasidharan et al 2018), we measure the posterior movement of the anterior tip of the pharynx. Retraction is not significantly altered in any of the 12 *hmr-1* mutant embryos assessed (Figure 2c), with an average of 17 microns involution in *hmr-1* mutants vs 18 microns for WT. This demonstrates that loss of neuroectoderm involution is independent of pharynx retraction in *hmr-1* mutant embryos and indicates additional function of *hmr-1* beyond apical constriction of the pharynx.

We then examine the effect of hmr-1 loss of function on the directed movement of individual neurons. To do so we measure the paths of a subset of neurons near the leading edge of the involution, namely SIBV, SIAD, AIY, and the mother of CEPV, on both the left and right side of the embryo based on live imaging and systematic cell lineage tracing (Bao et al 2006, Santella et al 2014, Katzman et al 2018). These neurons at the leading edge of the involuting neuroectoderm display directed movement towards the midline, while in *hmr-*1 mutants they no longer move towards the midline and instead move slightly anterior (Figure 2d). Convergent movement of left-right neurons towards each other is significantly reduced in *hmr-1* mutants in 3 out of 4 pairs measured, with between 40 and 70% less distance traveled in each (7.5-15 microns) (Figure 2e). These results reveal that the movement trajectories of neurons are shifted in *hmr-1* mutants.

### A Localized HMR-1 Patch at the Pharynx/Neuron Interface Mediates Intertissue Cohesion

A key aspect of our model is the attachment of neuroectoderm to the retracting pharynx. Given the *hmr-1* phenotypes, we ask if localized HMR-1 mediates this attachment (Figure 3a). Indeed, HMR-1::GFP (*xnIs96*, expressed under the control of *hmr-1* promoter), is strongly enriched at the interface between the basal pharyngeal surface and neuroectoderm during involution in a supracellular patch (Figure 3b, rectangle). This signal is distinct from the known localization at the apical side of the pharynx, which is dorsal to this patch (Figure 3b, arrowhead).

**Figure 3:**
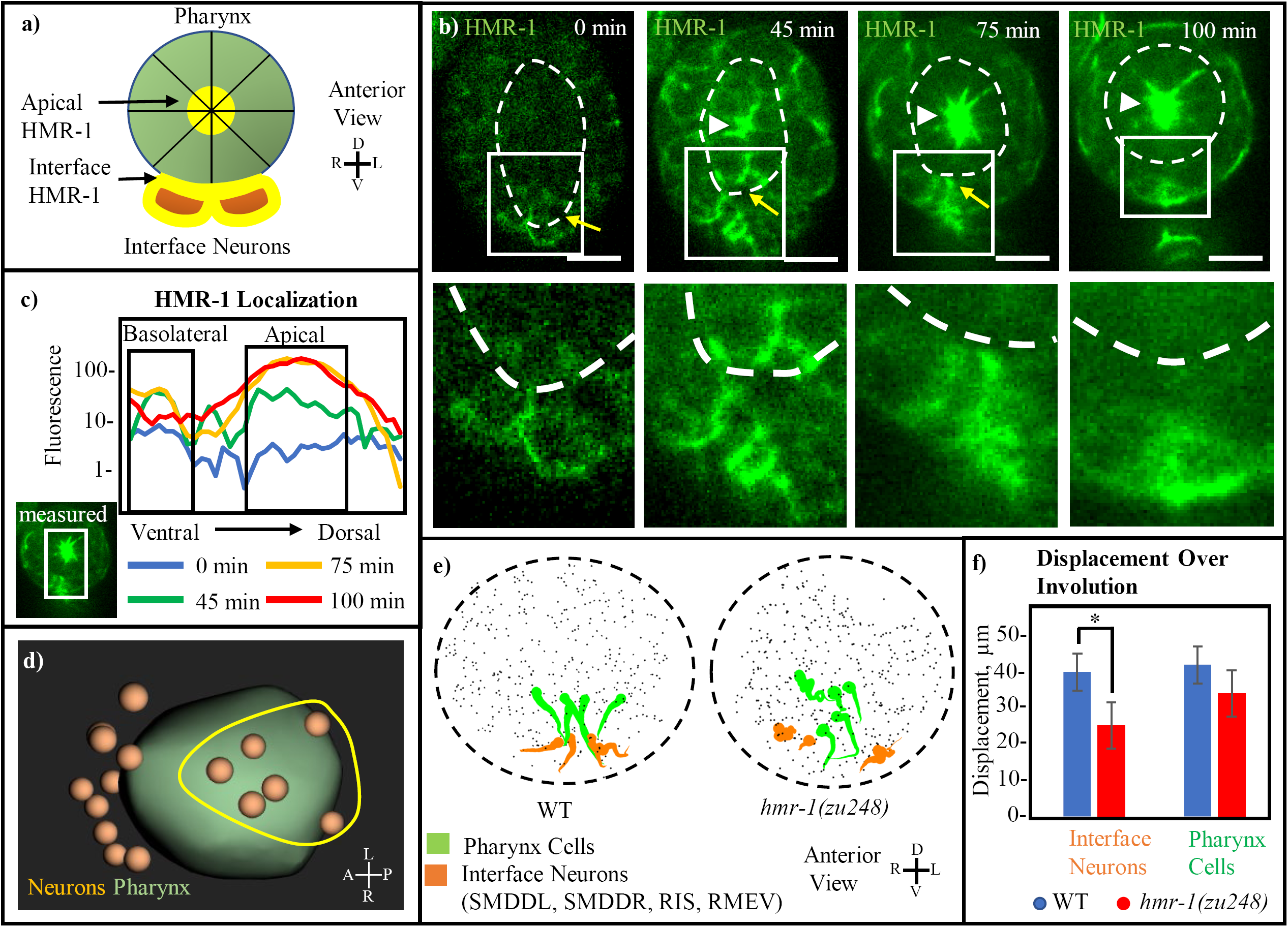
Cohesion at the Intertissue Interface is Maintained by a Local HMR-1 Patch. **(a)** Model of the adhesive HMR-1 interface (yellow) at the connection between ventral interface neurons (orange) and the retracting pharynx (green) mediating intertissue cohesion during involution. Apical HMR-1 is labelled, and lines through pharynx indicate apical constriction. **(b)** Anterior view HMR-1::GFP timeseries showing the basolateral (interface) and apical HMR-1 patches, starting before pharynx retraction when the interface patch is first forming, and ending after pharynx retraction when the interface patch disappears and only the apical localization remains. White arrowheads indicate apical HMR-1, yellow arrow indicates interface HMR-1. Cutouts show enlarged view of the interface at each timepoint. White dashed line indicates pharynx. Scale bars are 10µm. **(c)** Mean HMR-1::GFP fluorescence intensity at each plane from dorsal to ventral, analyzed in the 4 timepoints in e) and measured from the area illustrated. Basolateral and apical patches are indicated. **(d)** Interface neurons, defined as those with a persistent connection to a pharynx cell from the beginning of involution for at least 75% of timepoints, form a patch near the ventral midline (full data in Figure 3 Supplement 1 and Table 1). Yellow circle indicated rough shape of the group of interface neurons on the ventral side of the pharynx. **(e)** Anterior view shows motion paths of 4 interface neurons (orange) as well as their pharyngeal neighbors (green) in WT and *hmr-1(zu248)*. Tadpoles are as in 2e). Black dashed circle is embryo outline, grey dots are other cells. **(f)** Graph showing the total displacement over the timecourse of involution (n=3 embryos). Neurons: SMDDL, SMDDR, Pharynx Cells: Selected in each embryo according to proximity to SMDDL/R.

The existence of the HMR-1 patch at the neuron-pharynx interface is transient and coincides with the involution process (Figure 3 Movie 1). Prior to pharynx retraction, HMR-1 expression across the embryo is low, prior to pharynx retraction, but localization to the interface and the formation of the supracellular patch are evident (Figure 3b first image). Notably, this patch forms before there is meaningful signal of HMR-1 localization at the apical side of the pharynx. HMR-1::GFP signal rises and stays high during pharynx retraction and involution (Figure 3b second and third images), and disappears afterwards (Figure 3b fourth image, with the remaining signal in the rectangle belonging to skin cells). Quantification of HMR-1::GFP fluorescence intensity confirms this observation and further reveals the difference in temporal dynamics of HMR-1 localization between the supracellular patch at the neuron-pharynx interface and the apical side of the pharynx (Figure 3c), with the apical localization in the pharynx starting one step later and remaining high after involution completes.

Meanwhile, computational analysis of neighbor relationships between pharyngeal cells and the neurons adjacent to them further indicates where the attachment occurs at the physical interface. Specifically, we use Delaunay triangulation among nuclei to approximate neighbor relationship and detect cell pairs that persist as neighbors (see Methods, Figure 3 supplement 1a). Using data from 3 WT embryos where the entire cell lineage was traced up to the one-and-half fold stage (Santella et al 2015), we identify neurons that maintain their original pharyngeal neighbors during >75% of timepoints during involution (Figure 3 supplement 1b, Table 1). This analysis identified two groups of neurons, one on the anterior and one on the ventral side of the pharynx (Figure 3d). The ventral group (Figure 3d, yellow circle), which include SMDD, RIS and RMEV, spatially coincide with the supracellular HMR-1 patch, supporting the concept that HMR-1 at the basal pharyngeal surface promotes cohesion across the tissue interface. We are not able to examine HMR-1 localization at the anterior of the pharynx due to technical difficulties in orienting the tilting pharyngeal surface to the imaging axis.

**Table 1:** Persistence of pharynx contact across neurons and selected leader cells

| Cell | Timepoints<br>Contacting<br>Initial<br>Neighbors | %<br>Time | Cell | Timepoints<br>Contacting<br>Initial<br>Neighbors | %<br>Time | Cell | Timepoints<br>Contacting<br>Initial<br>Neighbors | %<br>Time |
| --- | --- | --- | --- | --- | --- | --- | --- | --- |
| 'AIAL' | 0 | 0.0 | 'IL2DR' | 27 | 41.5 | 'RMGR' | 0 | 0.0 |
| 'AIAR' | 0 | 0.0 | 'IL2L' | 0 | 0.0 | 'SAADL' | 0 | 0.0 |
| 'AIML' | 0 | 0.0 | 'IL2R' | 0 | 0.0 | 'SAADR' | 7 | 10.8 |
| 'AIMR' | 0 | 0.0 | 'IL2VL' | 0 | 0.0 | 'SAAVL' | 43 | 66.2 |
| 'AINL' | 35 | 53.8 | 'IL2VR' | 0 | 0.0 | 'SAAVR' | 18 | 27.7 |
| 'AINR' | 56 | 86.2 | 'OLLL' | 0 | 0.0 | 'SABD' | 0 | 0.0 |
| 'AIYL' | 39 | 60.0 | 'OLLR' | 0 | 0.0 | 'SABVL' | 0 | 0.0 |
| 'AIYR' | 40 | 61.5 | 'OLQDL' | 63 | 96.9 | 'SABVR' | 0 | 0.0 |
| 'ALA' | 0 | 0.0 | 'OLQDR' | 65 | 100.0 | 'SIADL' | 7 | 10.8 |
| 'AVAL' | 0 | 0.0 | 'OLQVL' | 0 | 0.0 | 'SIADR' | 23 | 35.4 |
| 'AVAR' | 0 | 0.0 | 'OLQVR' | 0 | 0.0 | 'SIAVL' | 51 | 78.5 |
| 'AVDL' | 65 | 100.0 | 'RIAL' | 0 | 0.0 | 'SIAVR' | 18 | 27.7 |
| 'AVDR' | 34 | 52.3 | 'RIAR' | 32 | 49.2 | 'SIBVL' | 0 | 0.0 |
| 'AVEL' | 0 | 0.0 | 'RID' | 16 | 24.6 | 'SIBVR' | 2 | 3.1 |
| 'AVER' | 0 | 0.0 | 'RIFL' | 0 | 0.0 | 'SMBDL' | 11 | 16.9 |
| 'AVG' | 51 | 78.5 | 'RIFR' | 0 | 0.0 | 'SMBDR' | 17 | 26.2 |
| 'AVHL' | 65 | 100.0 | 'RIGL' | 0 | 0.0 | 'SMBVL' | 0 | 0.0 |
| 'AVHR' | 54 | 83.1 | 'RIGR' | 0 | 0.0 | 'SMBVR' | 0 | 0.0 |
| 'AVJL' | 0 | 0.0 | 'RIH' | 27 | 41.5 | 'SMDDL' | 65 | 100.0 |
| 'AVJR' | 0 | 0.0 | 'RIPL' | 54 | 83.1 | 'SMDDR' | 53 | 81.5 |
| 'AVKL' | 37 | 56.9 | 'RIPR' | 0 | 0.0 | 'SMDVL' | 52 | 80.0 |
| 'AVKR' | 0 | 0.0 | 'RIR' | 41 | 63.1 | 'SMDVR' | 10 | 15.4 |
| 'AVL' | 33 | 50.8 | 'RIS' | 59 | 90.8 | 'URAVL' | 0 | 0.0 |
| 'BAGL' | 32 | 49.2 | 'RIVL' | 65 | 100.0 | 'URAVR' | 0 | 0.0 |
| 'BAGR' | 42 | 64.6 | 'RIVR' | 0 | 0.0 | 'URADL' | 0 | 0.0 |
| 'CEPDL' | 26 | 40.0 | 'RMDDL' | 22 | 33.8 | 'URADR' | 0 | 0.0 |
| 'CEPDR' | 31 | 47.7 | 'RMDDR' | 16 | 24.6 | 'URBL' | 0 | 0.0 |
| 'CEPVL' | 0 | 0.0 | 'RMDL' | 0 | 0.0 | 'URBR' | 0 | 0.0 |
| 'CEPVR' | 0 | 0.0 | 'RMDR' | 0 | 0.0 | 'URXL' | 26 | 40.0 |
| 'IL1DL' | 19 | 29.2 | 'RMDVL' | 0 | 0.0 | 'URXR' | 31 | 47.7 |
| 'IL1DR' | 19 | 29.2 | 'RMDVR' | 18 | 27.7 | 'URYDL' | 13 | 20.0 |
| 'IL1L' | 0 | 0.0 | 'RMED' | 16 | 24.6 | 'URYDR' | 44 | 67.7 |
| 'IL1R' | 42 | 64.6 | 'RMEL' | 41 | 63.1 | 'URYVL' | 0 | 0.0 |
| 'IL1VL' | 0 | 0.0 | 'RMER' | 54 | 83.1 | 'URYVR' | 0 | 0.0 |
| 'IL1VR' | 0 | 0.0 | 'RMEV' | 65 | 100.0 |  |  |  |
| 'IL2DL' | 36 | 55.4 | 'RMGL' | 0 | 0.0 |  |  |  |

We then examine if *hmr-1* is required for the correlated movement between the interface neurons and the adjacent pharyngeal cells. We measure this using the displacement of cells in WT and *hmr-1* mutant embryos (Figure 3e). Four of the interface cells, namely SMDDL, SMDDR, RIS and RMEV and the SMDD’s, show coordinated motion with their pharyngeal neighbors in WT embryos. In *hmr-1* mutants their movement trajectories diverge from their initial pharyngeal neighbors. These neurons move significantly less in *hmr-1* mutants compared to the WT, with an average of 38% (15 microns) less displacement measured across 3 embryos, while pharynx displacement is not significantly changed (Figure 3f). Based on the localization, phenotypes and computational analysis, we conclude that HMR-1 mediates the attachment between the interface neurons and the pharynx.

### Cohesion Within the Neuroectoderm Requires HMR-1

Furthermore, we examine whether HMR-1 is also required for cohesion within the neuroectoderm by examining its localization between neurons, and whether its loss would result in sliding between neighboring neurons (Figure 4a). HMR-1 localizes between neurons during involution, albeit weaker than at their interface with the pharynx (Figure 4b). To assay potential sliding, we examine the correlation of movement trajectory among neurons. Compared to the WT, neurons in *hmr-1* mutants show reduced, less directional movement towards the midline (Figure 4c). We further quantify the correlation between individual movement trajectories within a select group of neighboring neurons (SIBV, SIAD, AIY, and the mother of CEPV on the left and right side) (Figure 4d). In WT embryos, their respective movement is positively correlated on the ipsilateral side, and anticorrelated with the contralateral side. In contrast, *hmr-1* mutants show a general lack of correlation regardless of sides. This data is normalized across 3 WT and 3 mutant embryos. Together, the localization and phenotype assays support the hypothesis that HMR-1 also mediates intratissue cohesion within the neuroectoderm.

**Figure 4:**
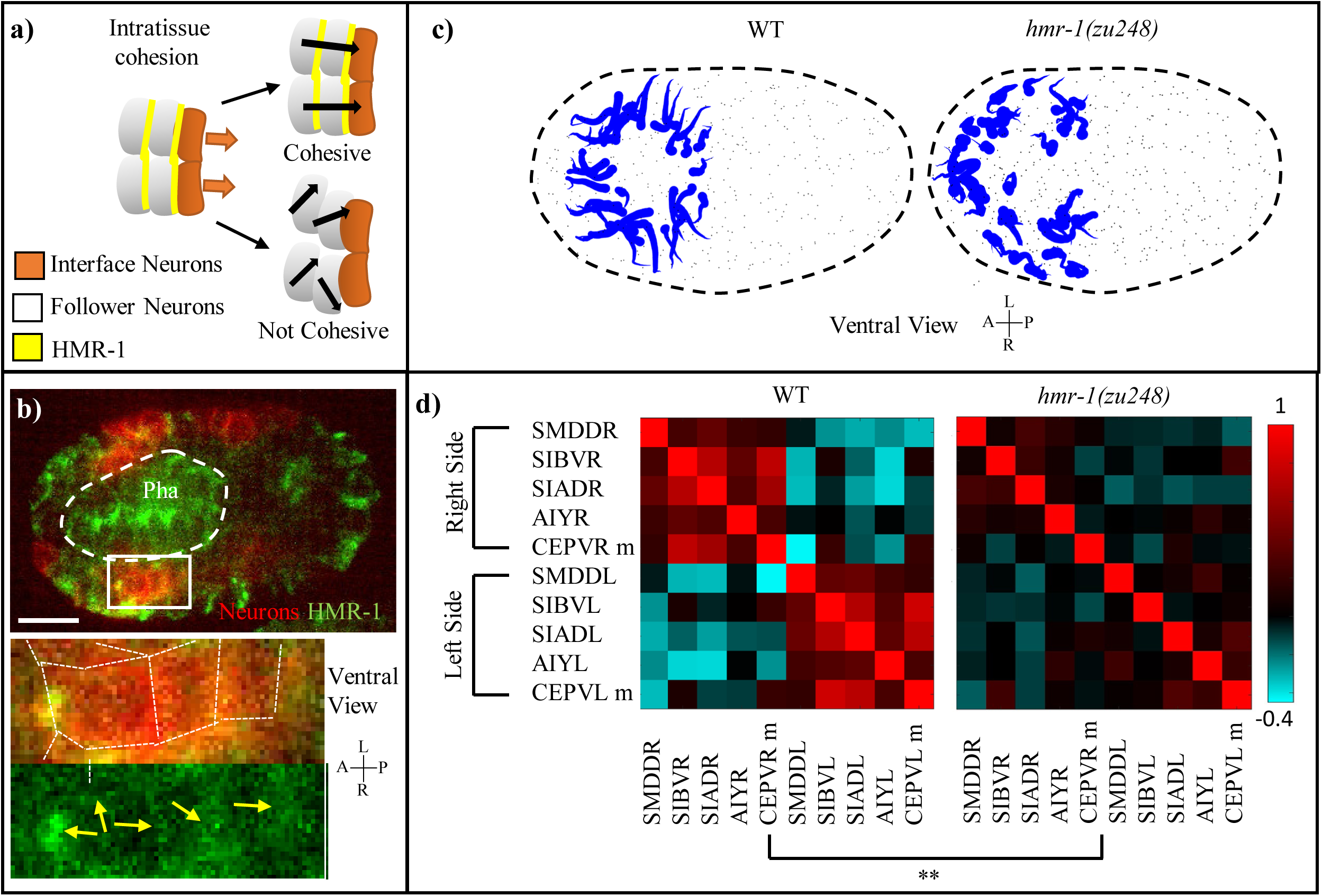
Intratissue Cohesion in the Neuroectoderm Also Requires *hmr-1*. **(a)** Model of expected neuronal behavior with and without intratissue cohesion. Yellow lines indicate adhesive interfaces between cells. Orange arrows mark the movement of interface neurons with the pharynx, black arrows mark the expected path of neurons. **(b)** HMR-1::GFP localization at the connection between involuting neurons. The patch of neurons is marked by *cnd1p*::PH::RFP expression and enlarged in cutout. White dashed line represents pharynx. Thin white dotted lines in cutout represent cell boundaries in patch of neurons, and yellow arrows indicate HMR-1::GFP signal at these cell boundaries. Scale bars = 10µm. **(c)** Motion paths of neurons in WT and *hmr-1(zu248)* embryos, tadpoles are as in 2e). Black dashed circle is embryo outline, grey dots are other cells. **(d)** Movement path correlation of select involuting neurons in a cluster on the ventral side, between WT (n=3) and *hmr-1(zu248)* (n=3) embryos. Bar indicates correlation value for each color (1=moving in same direction, −1 = moving in opposite directions). Left side vs right side neurons are labelled. P-value of .01 calculated with Paired T-test applied globally between neurons. M is mother.

### *C. elegans* Involution is Potentially Homologous to Vertebrate Neurulation

We find striking similarity between the coordinated tissue movement in *C. elegans* neuroectoderm involution versus vertebrate neurulation (Figure 5a). First, both have an apically constricting force generator; in *C. elegans* it is the pharynx and in vertebrates it is the floor plate/notochord (chordoneural hinge). In both cases the force generator is derived from outside of the neuroectoderm, and *pha-4/foxa2* is required in each case to specify the force generator (Horner et al 1998, Harrelson et al 2012) (Figure 5b). Hensen’s node (the vertebrate “organizer” region) is a *foxa2* specified tissue which proceeds to form three separate tissues across the germ layers (the floor plate, notochord, and dorsal endoderm) (Teillet et al 1998). The pharynx is derived from both the non-neuronal ectoderm lineage and the mesoderm. Second, topology and movement of the neurons, pharynx and skin is shared with vertebrates across the anterior-posterior axis (Figure 5a). The only difference is that involution and skin closure happen on the dorsal side in vertebrates and the ventral side in *C. elegans*. However, it is established that a flip of the D-V axis occurred in deuterostome evolution (Arendt & Nübler-Jung 1994). As discussed in more details below, these similarities may indicate evolutionary homology, and we propose that neuroectoderm involution in *C. elegans* should be considered as neurulation.

**Figure 5:**
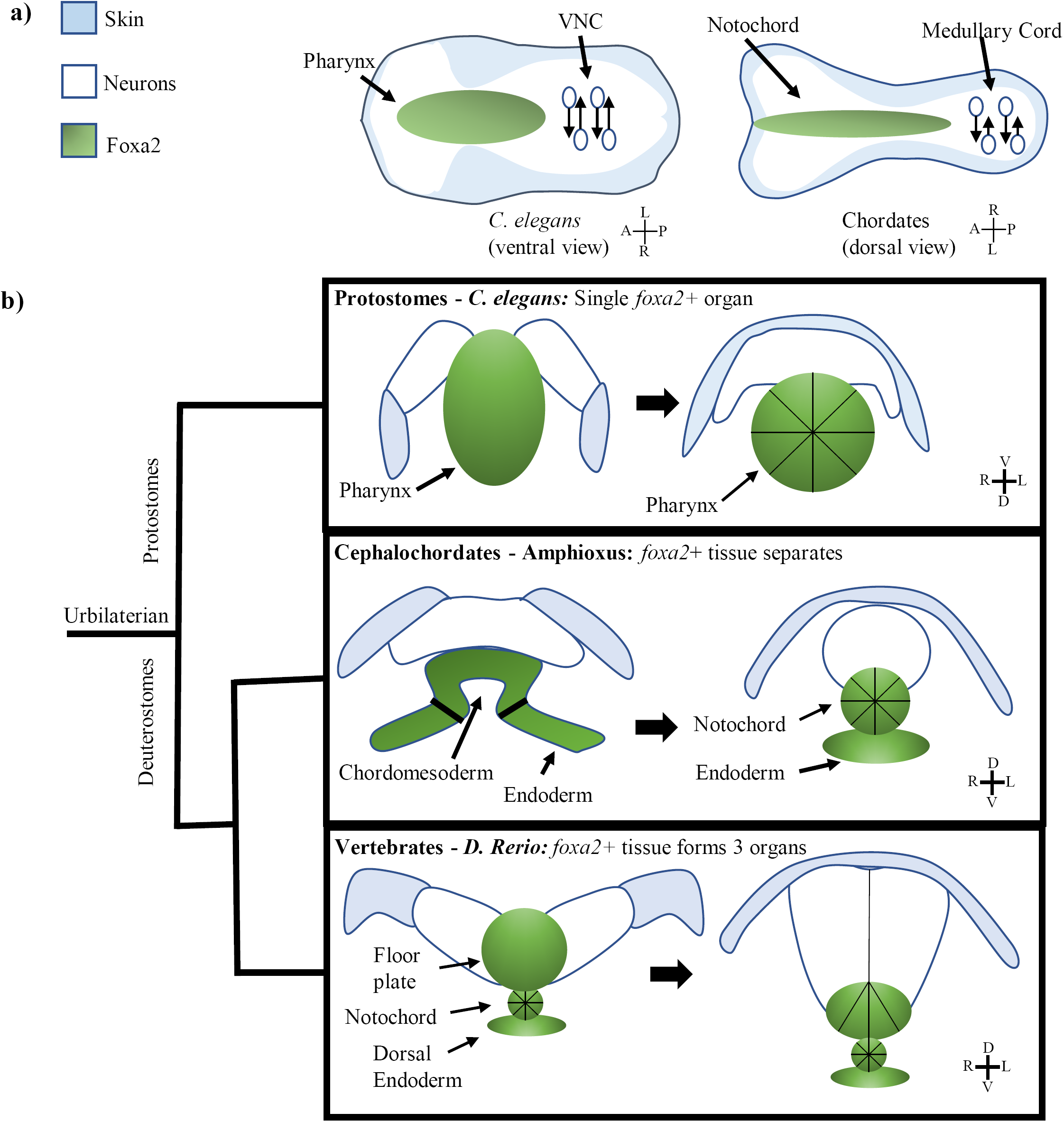
The Multi Tissue Process of *C. elegans* Involution May be Homologous to Chordate Neurulation. **(a)** Model of morphology during nervous system centralization in *C. elegans* as well as Chordates, labelling involution in the anterior of the embryo (pharynx in green, neuroectoderm as white background) and PCP/CE driven nerve cord formation in the posterior of the embryo (neurons as white circles). Hypoderm in in blue. Body axis is flipped between *C. elegans* and vertebrates (in b) as well). **(b)** Model showing foxa2 specified force generator during nervous system involution across 3 clades. Left images are before involution, right images are after. Black lines indicate apically constricting cells. Protostomes (*C. elegans)* have involution driven by a single foxa2+ tissue (the pharynx). Cephalochordates (Amphioxus) have involution driven by a foxa2+ layer which later forms 2 detached tissues (the notochord and gut). Chordates (*D. Rerio)* have separation of the foxa2+ tissue into 3 organs (floor plate, notochord, and dorsal endoderm) before involution.

## Discussion

### Role of Cadherins in *C. elegans* Nervous System Involution

Our work shows that loss of *hmr-1* in *C. elegans* has a similar phenotype to loss of N-cadherin (*cdh-2)* in zebrafish neurulation (Hong and Brewster 2006). In both cases, initial invagination and apical constriction in the foxa2+ tissue, namely the pharynx in *C. elegans* and the floor plate/notochord in zebrafish, is relatively unaffected (Hong and Brewster 2006), but involution of neuroectoderm fails. The role of E-cadherin (*cdh-1)* and N-cadherin in chordate neurulation has been heavily studied, with a shift from E-cadherin expression to N-cadherin expression specific to the neuroectoderm during this time period (Nandadasa et al 2012, Lele et al 2002). The *C. elegans hmr-1* is considered an orthologue of E-cadherin, but a splice variant is considered to act more similarly to N-cadherin (Broadbent and Pettitte 2002). Our perturbation did not separate the splice variant, though the phenotype is like that that of N-cadherin. More specific perturbations targeting the splice variants of *hmr-1*, such as CRISPR-based gene modification, may help resolve the issue.

Cadherins are well known for their established roles at the apical membrane during apical constriction, and for their presence on the lateral membrane to maintain cohesion within an epithelium. However, our work shows HMR-1 mediates adhesion on the basal side of the pharynx. If HMR-1 is in fact on the pharynx basal membrane it would raise an interesting question as to how a cell coordinates two distinct locations. On the other hand, if HMR-1 is only used in the interface neurons, it raises the question of what functions on the pharynx side to bind to HMR-1. Protocadherins, which are known to interact with N-cadherin in neurulation (Biswas et al 2010), are abundant in the *C. elegans* genome (Hardin et al 2013). It is worth noting that in vertebrates, functional attachment between the floor plate and the notochord (both of which are epithelial) occurs between their basal sides (Smith 1997). Upon closer examination of published data, N and E cadherin are both transiently localized to the basal side of the floor plate/neuroectoderm and notochord (Dady et al. 2012, figure 4)). Tissue-specific labeling and perturbation of HMR-1 will start to address these questions in *C. elegans* and should motivate similar efforts in vertebrates.

### Potential Evolutionary Homology and Implications

While neurons originated in radially symmetric animal phyla such as Ctenophora and Cnidaria, they began to assemble into complex centralized systems such as nerve cords and brains only in Bilateria. The traditional view holds that the centralization of the nervous system evolved multiple times, but this idea has been challenged recently. Multiple studies have suggested that the brain only evolved once in the ancestral bilaterian. These arguments are based on similarities in anatomy and gene expression between clades (Table 2).

**Table 2:** Previous articles supporting central nervous system developmental homology across Bilateria

| **Clade/Category** | **Papers** |
| --- | --- |
| Hemichordates | Miyamoto & Wada 2013 |
| Protostomes | Denes et al 2007, Hirth et al. 2003 |
| Acoelomorphs | Bailly et al 2013 |
| Cnidarians | Watanabe et al 2009 |
| Multiple Clades | Holland et al 2013 |
| *pha-4/foxa2*<br>expression | Arendt et al 2016, Magie et al 2005 |

The major argument against the idea that central nervous systems evolved once in an urbilaterian ancestor is that it only considers the molecular mechanism of lineage differentiation and neural fate specification. The shared transcription factors and cascades in neural fate specification are not directly involved in the centralization process and could have been recycled from an ancestral non-centralized nervous system. Therefore, to make a stronger argument for homology one should show among bilaterians shared functional cell biological aspects of nervous system centralization, and shared essential gene expression in tissues involved in coordinated morphogenesis beyond the nervous system itself. Since these aspects have been well characterized in vertebrates, which are deuterstomes, the strongest evidence would come from a protostome.

Our characterization of neuroectoderm involution in *C. elegans* is to our knowledge the first of its kind in protostomes, and the results satisfy the required arguments. Cnidarians have been shown to have a primitive nerve ring derived from neuroectoderm which initially ingresses a short distance (Watanabe et al 2009), and *D. Melanogaster* neuroectoderm moves to the midline after an involuting mesoderm layer (Mizutani and Bier 2010), but the underlying cell biological mechanism has not been characterized in either situation. In contrast, we have characterized the mechanism in depth and shown that, like in vertebrates, the neuroectoderm involution in *C. elegans* is driven by cohesive attachment to an apically constricting *pha-4/foxa2* specified tissue.

Comparative morphological analysis of clades phylogenetically in-between vertebrates and the ancestral bilaterian gives a potential explanation for traits derived in the deuterostome line. Amphioxus, which fits this phylogenetic requirement as a primitive chordate, also has involution of a neuroectoderm layer cohesively attached to a single apically constricting *pha-4/foxa2* specified tissue (Albuixech-Crespo et al 2017, Langeland et al 1999). More interestingly, Amphioxus presents a phenotypic halfway point between *C. elegans* and vertebrates; there is a single force generating layer attached to the neuroectoderm during involution (like in *C. elegans*) and the notochord separates from the force generator afterwards (Figure 5b). Hemichordates are deuterostomes which evolved earlier than amphioxus and also undergo neurulation. They have a partial notochord which never detaches from the endoderm (the “stomochord”) (Miyamoto and Wada 2013). These examples support an evolutionary model where the *foxa2+* tissue driving involution in each organism may have begun in the ancestral bilaterian as a single tissue attached to the neuroectoderm, and then gradually evolved in the deuterostome line to form a separate notochord and floor plate.

The role of *hmr-1/cadherin* in the processes of *C. elegans* involution and vertebrate neurulation further supports the concept of shared origins. *C. elegans* and zebrafish show similar phenotypes for loss of *hmr-1/cadherin*, and zebrafish may also use the non-canonical basal Cadherin to mediate intertissue attachment. While general use of Cadherin for adhesion can easily arise through convergent evolution, a shared non-canonical role at the basal side of an epithelium is perhaps less likely to do so.

Lastly, the morphogenesis of the VNC of *C. elegans*, as characterized in our previous study, also draws significant parallels to vertebrate neurulation. Both processes involve PCP mediated cell intercalation and convergent extension to elongate the nervous system along the anteroposterior axis, (Williams et al 2014, Shah et al 2017), thus highlighting an additional key morphological mechanism of neurulation shared between a protostome and vertebrates. Overall, our findings provide a strong argument from the perspective of shared developmental morphology and a conserved force generator to favor the hypothesis that the brain only evolved once.

## Materials and Methods

### *C. elegans* Strains and Genetics

*C. elegans* strains were grown on NGM plates seeded with OP50 bacteria as previously detailed (Brenner 1974). N2 Bristol was used as the WT strain. All worms were grown at room temperature. The following strains were used throughout the study: *BV292(zyIs36[cnd-1p::PH::RFP] IV), BV308 (zyIs36 [cnd1-p::PH::RFP]X, unc-119(ed3) III; xnIs96 [hmr-1p::hmr-1::GFP::unc-54 3’UTR + unc-119(+)]), BV729 (zbIs12 [Punc-33_PHD_GFP_unc54 UTR, Punc-122_RFP]; ujIs113), BV731 (zbIs12; ujIs113, hmr-1(zu248) I; zuEx2), DCR4318 (olaex2540 [Punc-33_PHD_GFP_unc54, Punc-122_RFP]; ujIs113), BV745(zyIs36[cnd-1p::PH::RFP] IV, hmr-1(zu248) I; zuEx2)*.

### Embryonic Imaging

Preparation of embryos for live imaging was done as previously described (Bao and Murray 2011). Gravid adult worms were picked to ∼30 µl M9 buffer (3 g KH2PO4, 6 g Na2HPO4, 5 g NaCl, 1 ml 1 M MgSO4, per liter H2O), transferred to a second drop to dilute extra OP50 bacteria, and cut open to release embryos. Depending on the goal of the imaging experiment, one of three next steps would be taken. When embryos were to have their lineage analyzed using AceTree software, 4-10 embryos at 2-4 cell stage were transferred to a small drop (∼1.5µl) of M9 media mixed with 20 µm polystyrene beads on a 24 x 50mm coverslip in and sealed with Vaseline under an 18 x 18mm smaller coverslip. When embryos were intended only for visual phenotyping, the same protocol was used, but up to 40 embryos were imaged at once. Lastly, when embryos were to be viewed through an anterior-posterior orientation, embryos were added to a larger (∼4µl) drop of M9 without beads, a thin layer of Vaseline was deposited above and below the drop, and the 18 x 18mm coverslip was added on top and further sealed with an extra layer of Vaseline to allow uncompressed imaging.

Images were acquired on either a spinning disk confocal microscope comprising a Zeiss Axio Observer Z1 frame with an Olympus UPLSAPO 60XS objective, a Yokogawa CSU-X1 spinning-disk unit, and two Hamamatsu C9100-13 EM-CCD cameras, or an instant structured illumination microscope (Visitech iSIM) using an Olympus IX73 body, an Olympus UPLSAPO40XS objective and a Hamamatsu Flash 4.0v2 sCMOS camera. Z stacks composing 30 slices of 1 micron each were used for imaging of compressed embryos, and 36 slices of 2 microns each for uncompressed embryos. Embryos were imaged every 75 seconds (lineaging) or between 2-5 minutes (non-lineaging), sufficient time was allowed to enable visualization of terminal phenotypes for easier selection of mutants. Imaging exposure time was 150ms per slice, with 568nm laser exposure every timepoint and 488nm laser exposure every 5 minutes regardless of imaging frequency.

### Image Analysis

Visual analysis of embryos was done using Fiji software (Schindelin et al 2012). In figure 1, Visual tracking of involuting chains was done with Fiji. In figure 2, to determine degree of involution we measured the distance between the leading edge of involuting neurons as visible with cnd1p::RFP marker on Fiji. To measure degree of pharynx retraction, distance to the anterior tip of the fully retracted pharynx from the anterior tip of the embryo was also measured manually on Fiji. In figure 3 HMR-1 localization measurements, HMR-1:GFP fluorescence was measured starting immediately ventral of the pharynx dorsally past the apical side, taking the maximum across the width of the inner pharynx and subtracting baseline fluorescence.

### Computational Cell Motion Analysis

For automated tracking and rendering of cell movement paths, StarryNite software was used to segment RFP tagged nuclei (Santella et al 2010) and AceTree was used to edit cells of interest to assure successful tracking (Boyle et al 2006). All visualization and analysis done on single ID’d cells was done using MATLAB software, including renderings of cell movement patterns, cell displacement analysis, and motion path correlation analysis.

Cell Motion: Nuclear positions for each desired cell were extracted over the selected time window. When cell divisions occurred during this window desired cells and their parents were considered equivalent and their positions concatenated. Cell position over time was smoothed using the MATLAB smoothdata function which computes a moving window average at an automatically selected scale (see Figure 2d and e, Figure 3g and h, Figure 4c and d).

Correlation Analysis: Positions were differenced and correlation of 3d directional velocity over all timepoints was computed for all pairs of cells (see Figure 4d)

### Cohesive Neuron Screen

A screen was performed for neurons that maintain close contact with the pharynx over time. Based on Nuclear positions a Voronoi diagram approximation of cell-cell contacts was computed. The total of neuron-pharynx contacts over time was computed from this model, using only pharyngeal neighbors from the first timepoint. A threshold of 75% indicating a neuron with near constant contact with at least 1 pharynx cell was established and used to determine a set of leader neurons. The SMDD neurons, the most highly cohesive left/right neuron pair in the ventral patch of neurons, were selected for use in further computational analysis.

### WormGUIDES analysis

WormGUIDES is a tool which enables visualization of fine spatiotemporal analysis of selected cell movement patterns in the *C. elegans* embryo via an adjustable 3D rendering (Santella et al 2015). Involuting neuronal nuclei were visualized next to pharynx and hypodermal surface models, including visualizing the movement of these features overtime created using the FIJI temporal max projection tool.

## Supporting information

Key Resources Table

## Acknowledgements

We thank Dr. Shyr-Shea Chang and Braden Katzman for assistance with experiments and quantification, and all Bao lab members for general help and feedback. We thank Drs. Hari Shroff, Jeremy Dittman, Pavak Shah and Srivarsha Rajshekar for comments on the manuscript. Some strains were provided by the CGC, which is funded by NIH (P40 OD010440). This work was partly supported by NIH grants (R01 GM097576 and R24 OD016474) to Z.B. and a Core Grant to MSKCC (P30 CA008748). Research in the D.A.C.-R. lab was supported by NIH grant No. R24-OD016474 and by an HHMI Scholar Award. M.W.M was supported by NIH by F32-NS098616. A.S. was supported by grant 2019-198110 (5022) from the Chan Zuckerberg Initiative and the Silicon Valley Community Foundation.

---

**Figure 1 Supplement 1:**
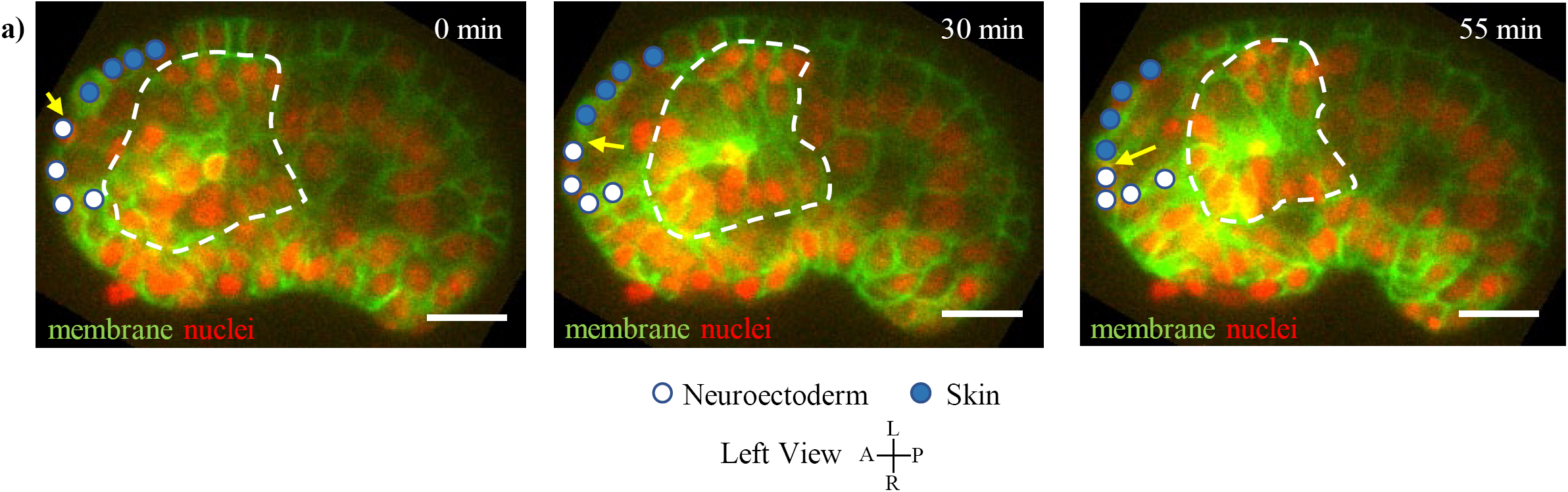
Persistent Contact Between Involuting Neuroectoderm and Skin. **(a)** Hypodermal cells (marked by blue circles) and neuroectodermal cells (marked by white circles) during involution, shown in *unc33p::PH::GFP* expressing embryos (left view, pharynx marked by white dashed circle). Yellow arrow indicates interface between hypodermal cells and neuroectoderm. Red channel is cell nuclei. Scale bars are 10µm.

**Figure 3 Supplement 1:**
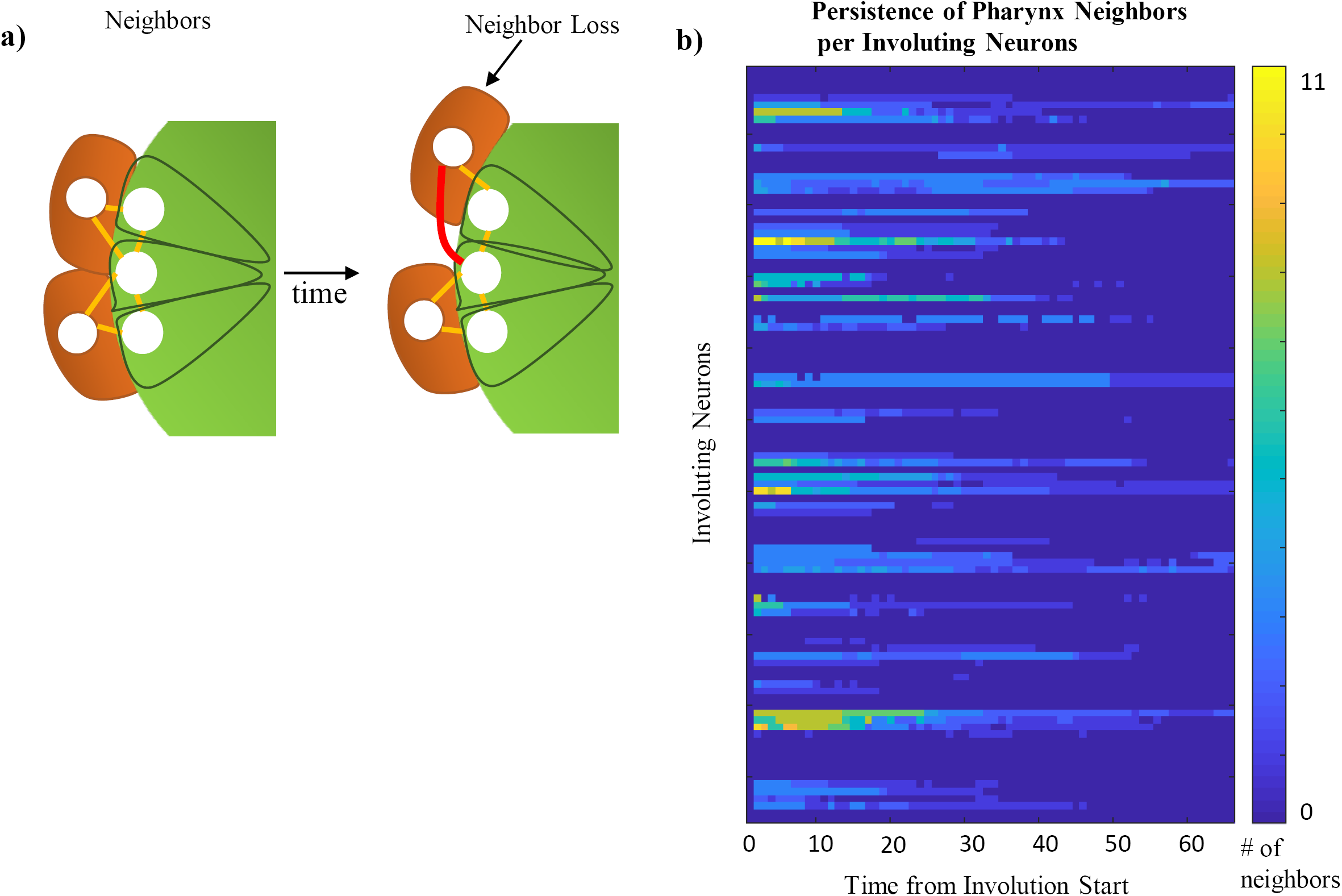
Computational Assessment of Persistent Cohesion from Relative Nuclear Location. **(a)** Model of a Delauney triangulation of an optimized cell nucleus-cell nucleus Voronoi graph. If an edge between two cells cannot be drawn without intersecting the nucleus of another cell (red line), the cells are not neighbors. If it can (orange lines), they are neighbors **(b)** Quantity of initial pharyngeal neighbors maintained over time for each neuron as measured by Delauney Triangulation. Coloring of blue to yellow indicates total # of pharyngeal neighbors to a given neuron at any timepoint. Full data in Table 1.

