## Supplementary Data for "Cadherin Preserves Cohesion Across Involuting Tissues During *C. elegans* Neurulation"

### Supplementary Data: List of Neurons Used in Figure Panels

| | Fig 1a, b | Fig 4c | | Fig 1a, b | Fig 4c | | Fig 1a, b | Fig 4c |
|---|---|---|---|---|---|---|---|---|
| AIAL | x | | IL2DL | x | | RMGR | x | |
| AIAR | x | | IL2DR | x | | SAADL | x | |
| AIML | x | X | IL2L | x | | SAADR | x | |
| AIMR | x | X | IL2R | x | | SAAVL | x | x |
| AINL | x | X | IL2VL | x | | SAAVR | x | x |
| AINR | x | X | IL2VR | x | | SABDL | x | |
| AIYL | x | X | OLLL | x | | SABDR | x | |
| AIYR | x | X | OLLR | x | | SABVL | x | |
| ALA | x | X | OLQDL | x | | SABVR | x | |
| AVAL | x | X | OLQDR | x | | SIADL | x | x |
| AVAR | x | X | OLQVL | x | | SIADR | x | x |
| AVDL | x | X | OLQVR | x | | SIAVL | x | |
| AVDR | x | X | RIAL | x | | SIAVR | x | |
| AVEL | x | | RIAR | x | | SIBVL | x | x |
| AVER | x | | RID | x | | SIBVR | x | x |
| AVG | x | | RIFL | x | | SMBDL | x | |
| AVHL | x | X | RIFR | x | | SMBDR | x | |
| AVHR | x | X | RIGL | x | | SMBVL | x | |
| AVJL | x | | RIGR | x | | SMBVR | x | |
| AVJR | x | | RIH | x | | SMDDL | x | x |
| AVKL | x | | RIPL | x | X | SMDDR | x | x |
| AVKR | x | | RIPR | x | X | SMDVL | x | x |
| AVL | x | X | RIR | x | | SMDVR | x | x |
| BAGL | x | | RIS | x | | URAVL | x | x |
| BAGR | x | | RIVL | x | X | URAVR | x | x |
| CEMVL | x | x | RIVR | x | X | URADL | x | |
| CEMVR | x | x | RMDDL | x | | URADR | x | |
| CEPDL | x | | RMDDR | x | | URBL | x | |
| CEPDR | x | | RMDL | x | | URBR | x | |
| CEPVL | x | | RMDR | x | x | URXL | x | |
| CEPVR | x | | RMDVL | x | x | URXR | x | |
| IL1DL | x | | RMDVR | x | | URYDL | x | |
| IL1DR | x | | RMED | x | x | URYDR | x | |
| IL1L | x | | RMEL | x | x | URYVL | x | x |
| IL1R | x | | RMER | x | x | URYVR | x | x |
| IL1VL | x | | RMEV | x | | | | |
| IL1VR | x | | RMGL | x | | | | |
